# Gut Microbiota Diversity across Ethnicities in the United States

**DOI:** 10.1101/342915

**Authors:** Andrew W. Brooks, Sambhawa Priya, Ran Blekhman, Seth R. Bordenstein

## Abstract

Composed of hundreds of microbial species, the composition of the human gut microbiota can vary with chronic diseases underlying health disparities that disproportionally affect ethnic minorities. However, the influence of ethnicity on the gut microbiota remains largely unexplored and lacks reproducible generalizations across studies. By distilling associations between ethnicity and differences in two United States based 16S gut microbiota datasets including 1,673 individuals, we report 12 microbial genera and families that reproducibly vary by ethnicity. Interestingly, a majority of these microbial taxa, including the most heritable bacterial family, Christensenellaceae, overlap with genetically-associated taxa and form co-occurring clusters linked by similar fermentative and methanogenic metabolic processes. These results demonstrate recurrent associations between specific taxa in the gut microbiota and ethnicity, providing hypotheses for examining specific members of the gut microbiota as mediators of health disparities.

## Introduction

The human gut microbiota at fine resolution varies extensively between individuals (1–3), and this variability frequently associates with diet (4–7), age (6, 8, 9), sex (6, 9, 10), body mass index (BMI) (1, 6), and diseases presenting as health disparities (11–14). The overlapping risk factors and burden of many chronic diseases disproportionally affect ethnic minorities in the United States, yet the underlying biological mechanisms mediating these substantial disparities largely remain unexplained. Recent evidence is consistent with the hypothesis that ethnicity associates with variation in microbial abundance, specifically in the oral cavity, gut, and vagina (15–17). To varying degrees, ethnicity can capture many facets of biological variation including social, economic and cultural variation, as well as aspects of human genetic variation and biogeographical ancestry. Ethnicity also serves as a proxy to characterize health disparity incidence in the United States, and while factors such as genetic admixture create ambiguity of modern ethnic identity, self-declared ethnicity has proven a useful proxy for genetic and socioeconomic variation in population scale analyses, including in the Human Microbiome Project (18–20). Microbiota differences have been documented across populations that differ in ethnicity as well as in geography, lifestyle, and sociocultural structure; however, these global examinations cannot disconnect factors such as intercontinental divides and hunter-gatherer versus western lifestyles from ethnically structured differences (21–23). Despite the importance of understanding the interconnections between ethnicity, microbiota, and health disparities, there are no reproducible findings about the influence of ethnicity on differences in the gut microbiota and specific microbial taxa in diverse United States populations, even for healthy individuals (6).

Here, we comprehensively examine connections between self-declared ethnicity and gut microbiota differences across more than a thousand individuals sampled by the American Gut Project (AGP, N=1375) (24) and the Human Microbiome Project (HMP, N=298) (6). Previous studies demonstrated that human genetic diversity in the HMP associates with differences in microbiota composition(25), and genetic population structure within the HMP generally delineates self-declared ethnicity (20). Ethnicity was not found to have a significant association with microbiota composition in a Middle Eastern population, however factors such as lifestyle and environment that influence microbiota variation across participants was homogenous compared to the ethnic, sociocultural, economic, and dietary diversity found within the United States (26). While ethnic diversity is generally underrepresented in current microbiota studies, evidence supporting an ethnic influence on microbiota composition among first generation immigrants has been recently demonstrated in a Dutch population (27). The goal of this examination is to evaluate, for the first time, if there are reproducible differences in gut microbiota across ethnicities within an overlapping United States population, as ethnicity is one of the key defining factors for health disparity incidence in the United States. Lifestyle, dietary, and genetic factors all vary to different degrees across ethnic groups in the United States, and it will require more even sampling of ethnic diversity and stricter phenotyping of study populations to disentangle which factors underlie ethnic microbiota variation in the AGP and HMP.

## Results

### Microbiota are subtly demarcated by ethnicity

We first evaluate gut microbiota distinguishability between AGP ethnicities (**Fig 1A**, family taxonomic level, Asians-Pacific Islanders (N=88), Caucasians (N=1237), Hispanics (N=37), and African Americans (N=13)), sexes (female (N=657), male (N=718)), age groups (years grouped by decade), and categorical BMI (underweight (N=70), normal (N=873), overweight (N=318), and obese (N=114)) (Demographic details in **S1A Table**). Age, sex, and BMI were selected as covariates because they are consistent across the AGP and HMP datasets. Additionally, 31 other categorical factors measuring diet, environment, and geography were compared for pairwise differences between two ethnicities using proportions tests, and very few (10 / 894) tests significantly varied (**S1 Table** additional sheets). Interindividual gut microbiota heterogeneity clearly dominates; however, Analyses of Similarity (ANOSIM) reveal subtle but significant degrees of total microbiota distinguishability for ethnicity, BMI, and sex, but not for age (**Fig 1B**, Ethnicity; **Fig 1C**, BMI; **Fig 1D**, Sex; **Fig 1E**, Age) (28). Recognizing that subtle microbiota distinguishability between ethnicities may be spurious, we independently replicate the ANOSIM results from HMP African Americans (N=10), Asians (N=34), Caucasians (N=211) and Hispanics (N=43) (**S2A Table**, R=0.065, p=0.044). We again observe no significant distinguishability for BMI, sex, and age in the HMP. Higher rarefaction depths increase microbiota distinguishability in the AGP across various beta diversity metrics and categorical factors (**S2B Table**), and significance increases when individuals from overrepresented ethnicities are subsampled from the average beta diversity distance matrix (**S2C Table**). Supporting the ANOSIM results, Permutational Multivariate Analysis of Variance (PERMANOVA) models with four different beta diversity metrics showed that while all factors had subtle but significant associations with microbiota variation when combined in a single model, effect sizes were highest for ethnicity in 7 out of 8 comparisons across beta diversity metrics and rarefaction depths in the AGP and HMP (**S2D Table**). We additionally test microbiota distinguishability by measuring the correlation between beta diversity and ethnicity, BMI, sex, and age with an adapted BioEnv test (**S2E Table**) (29). Similar degrees of microbiota structuring occur when all factors are incorporated (Spearman Rho=0.055, p-values: Ethnicity=0.057, BMI<0.001, Sex<0.001, Age=0.564). Firmicutes and Bacteroidetes dominated the relative phylum abundance, with each representing between 35% and 54% of the total microbiota across ethnicities (**S1 Fig**).

**Fig 1.**
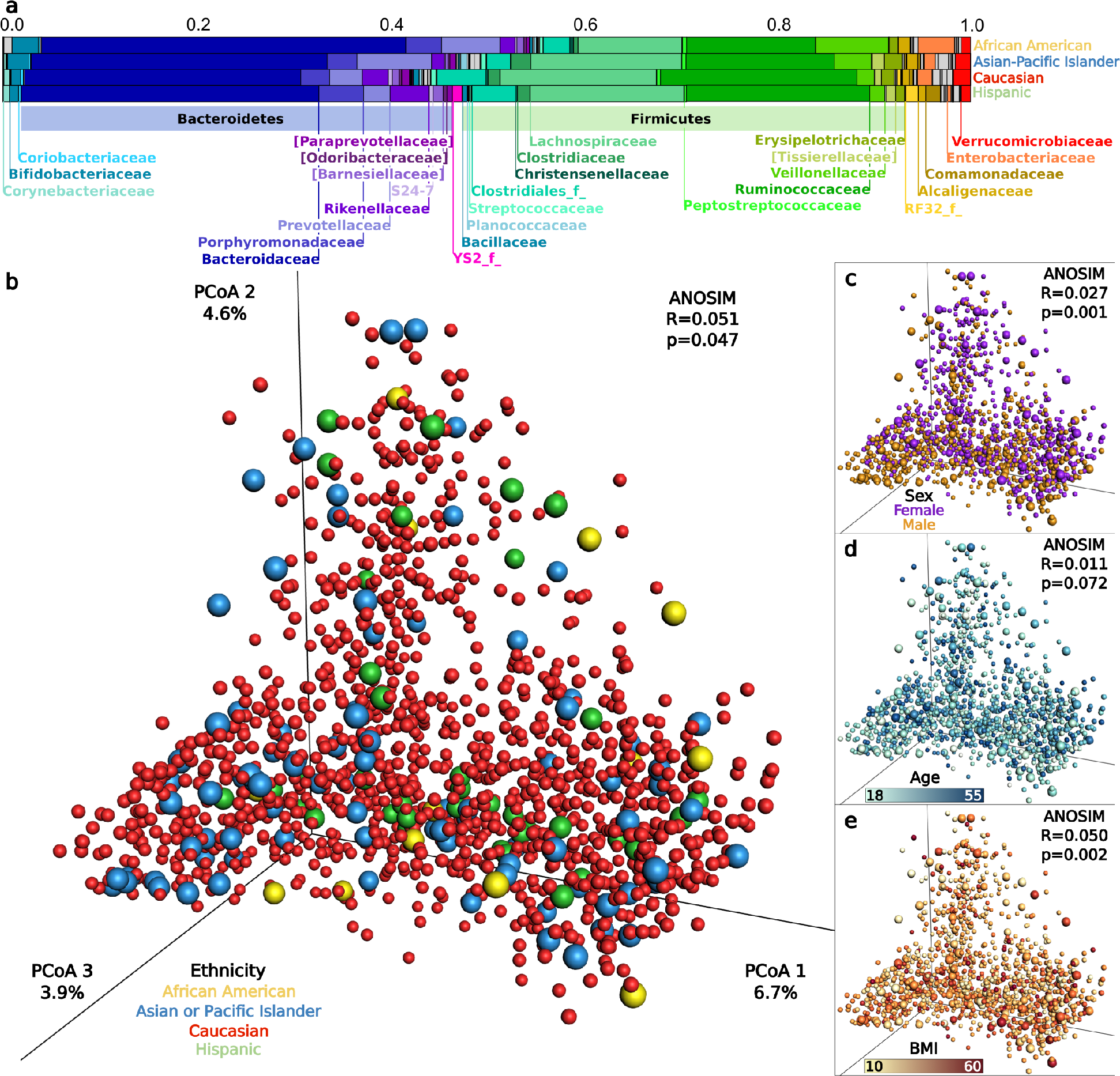
Gut microbiota composition and distinguishability by ethnicity, sex, age and BMI. (A) The average relative abundance of dominant microbial families for each ethnicity. (B-E) Principle coordinates analysis plots of microbiota Bray-Curtis beta diversity and ANOSIM distinguishability for: (B) Ethnicity, (C) Sex, (D) Age, (E) BMI. In B-E, each point represents the microbiota of a single sample, and colors reflect metadata for that sample. Caucasian points are reduced in size to allow clearer visualization, and p-values are not corrected across factors which have different underlying population distributions.

We next test for ethnicity signatures in the gut microbiota by analyzing alpha and beta diversity, abundance and ubiquity distributions, distinguishability, and classification accuracy (30). Shannon’s Alpha Diversity Index (31), which weights both microbial community richness (Observed OTUs) and evenness (Equitability), significantly varies across ethnicities in the AGP dataset (Kruskal Wallis, p=2.8e-8) with the following ranks: Hispanics > Caucasians > Asian-Pacific Islanders > African Americans (**Fig 2A**). In the HMP, there is a significantly lower Shannon diversity for Asian-Pacific Islanders relative to Caucasians and a trend of lower Shannon diversity for Asian-Pacific Islanders relative to Hispanics; African Americans change position in diversity relative to other ethnicities, potentially as a result of undersampling bias. Five alpha diversity metrics, two rarefaction depths, and separate analyses of Observed OTUs and Equitability generally confirm the results (**S3A Table**).

**Fig 2.**
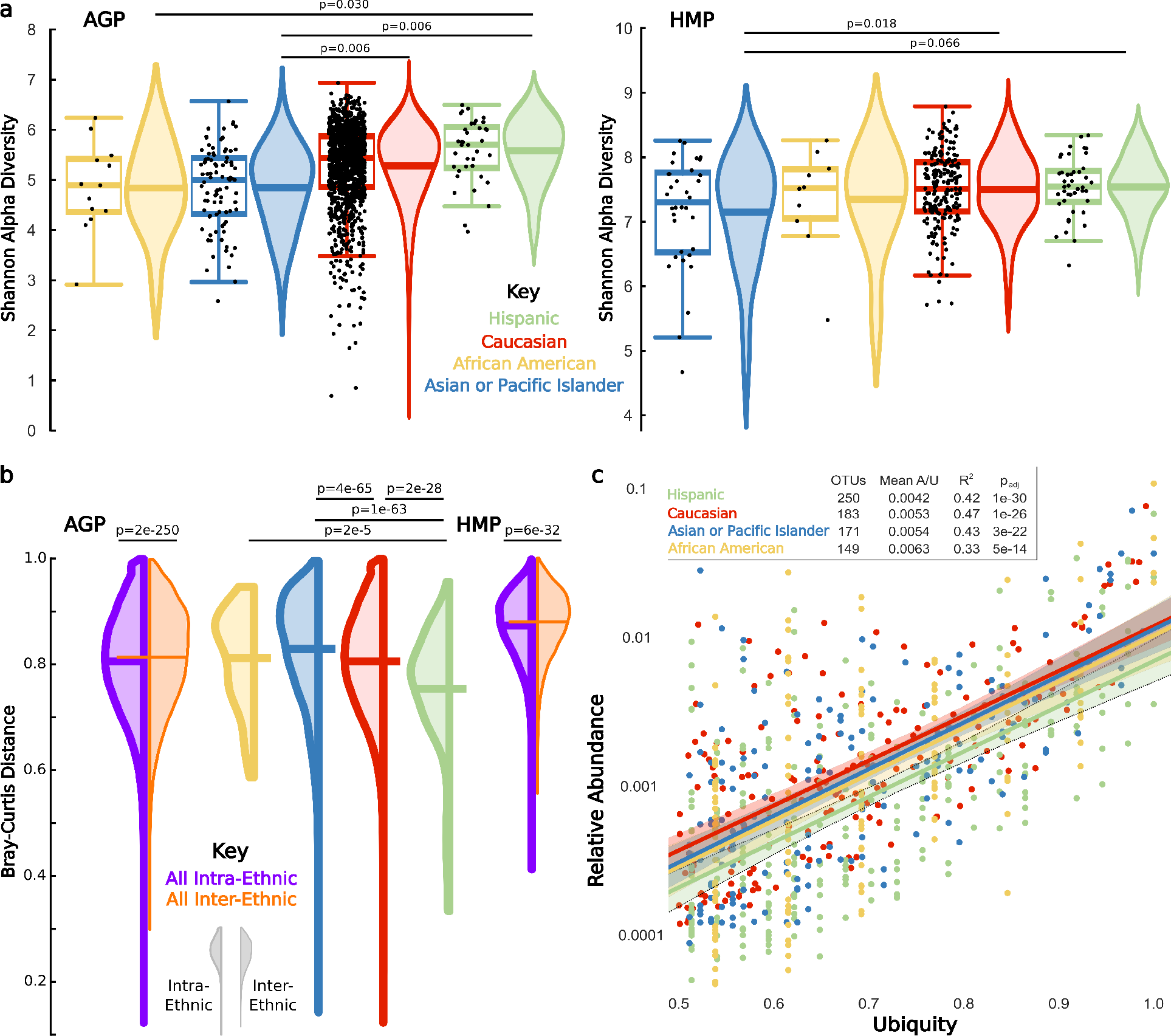
Ethnicity associates with diversity and composition of the gut microbiota. (A) Center lines of each boxplot depict the median by which ethnicities were ranked from low (left) to high (right); the lower and upper ends of each box represent the 25th and 75th quartiles respectively; whiskers denote the 1.5 interquartile range, and black dots represent individual samples. Lines in the middle of violin plots depict the mean, and p-values are Bonferroni corrected within each dataset. (B) Left extending violin plots represent intra-ethnic distances for each ethnicity, and right extending violin plots depict all inter-ethnic distances. Center lines depict the mean beta diversity. Significance bars above violin plots depict Bonferroni corrected pairwise Mann-Whitney-U comparisons of the intra-intra- and intra-inter-ethnic distances. (C) Within each ethnicity, OTUs shared by at least 50% of samples. Colored lines represent a robust ordinary least squares regression within OTUs of each ethnicity, shaded regions represent the 95% confidence interval, R^2^ denotes the regression correlation, the OTUs column indicates the number of OTUs with >50% ubiquity for that ethnicity, Mean A/U is the average abundance/ubiquity ratio, and the p_adj_ is the regression significance adjusted and Bonferroni corrected for the number of ethnicities.

If ethnicity impacts microbiota composition, pairwise beta diversity distances (ranging from 0/completely dissimilar to 1/identical) will be greater between ethnicities than within ethnicities. While average gut microbiota beta diversities across all individuals are high (**Fig 2B**, Bray-Curtis=0.808), beta diversities between individuals of the same ethnicity (intra-ethnic, Bray-Curtis=0.806) are subtly, but significantly, lower than those between ethnicities in both the AGP (inter-ethnic, Bray-Curtis=0.814) and HMP datasets (intra-ethnic, Bray-Curtis=0.870 versus inter-ethnic, Bray-Curtis=0.877). We confirm AGP results by subsampling individuals from overrepresented ethnicities across beta metrics and rarefaction depths (**S4A-4B Tables**). Finally, we repeat analyses across beta metrics and rarefaction depths using only the average distance of each individual to all individuals from the ethnicity to which they are compared (**S4C-4D Tables**).

Next, we explore inter-ethnic differences in the number of OTUs shared in at least 50% of individuals within an ethnicity, as the likelihood of detecting a biological signal is improved in more abundant organisms relative to noise that may predominate in lower abundance OTUs. Out of 5,591 OTUs in the total AGP dataset, 101 (1.8%) meet this ubiquity cutoff in all ethnicities, and 293 (5.2%) OTUs meet the cutoff within at least one ethnicity. Hispanics share the most ubiquitous OTUs and have the lowest average abundance/ubiquity (A/U) ratio (**Fig 2C**), indicating stability whereby stability represents a more consistent appearance of OTUs with lower abundance but higher ubiquity (32). This result potentially explains their significantly lower intra-ethnic beta diversity distance and thus higher microbial community overlap relative to the other ethnicities (**Fig 2B**). Comparisons in the AGP between the higher sampled Hispanic, Caucasian, and Asian-Pacific Islander ethnicities also reveal a trend wherein higher intra-ethnic community overlap (**Fig 2B**) parallels higher numbers of ubiquitous OTUs (**Fig 2C**), higher Shannon Alpha diversity (**Fig 2A**), and higher stability of ubiquitous OTUs as measured by the abundance/ubiquity (A/U) ratio (**Fig 2C**).

We next assess whether a single ethnicity disproportionately impacts total gut microbiota distinguishability in the AGP by comparing ANOSIM results from the consensus beta diversity distance matrix when each ethnicity is sequentially removed from the analysis (**Fig 3A** and **S2E Table**). Distinguishability remains unchanged when the few African Americans are removed, but is lost upon removal of Asian-Pacific Islanders or Caucasians, likely reflecting their higher beta diversity distance from other ethnicities (**Fig 3A**). Notably, removal of Hispanics increases distinguishability among the remaining ethnicities, which may be due to higher degree of beta diversity overlap observed between Hispanics and other ethnicities (**S4B Table**). Results conform across rarefaction depths and beta diversity metrics (**S2F Table**), and pairwise combinations show strong distinguishability between African Americans and Hispanics (ANOSIM, R=0.234, p=0.005), and Asian-Pacific Islanders and Caucasians (ANOSIM, R=0.157, p<0.001).

**Fig 3.**
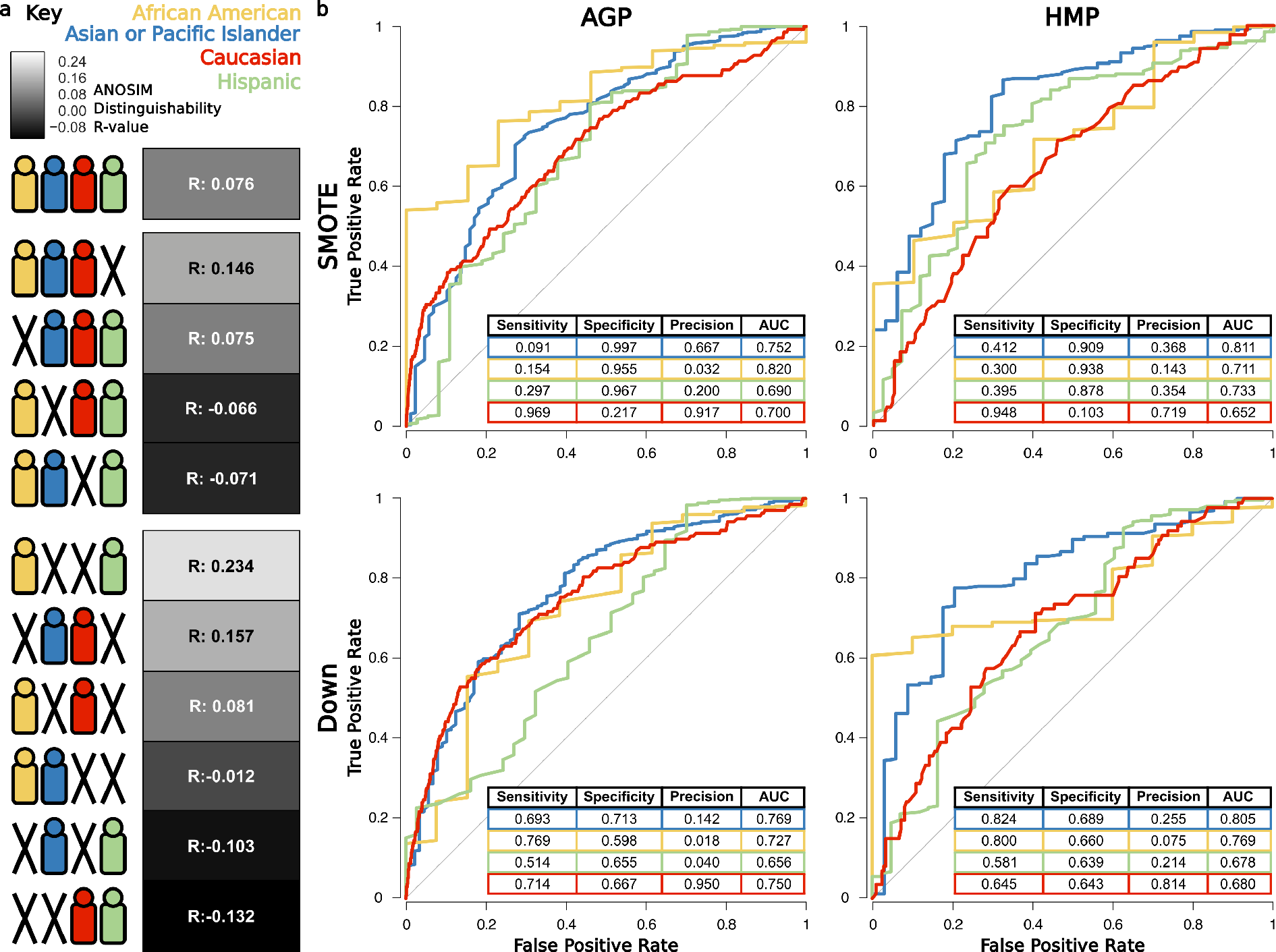
Microbiota distinguishability and classification ability across ethnicities. (A) ANOSIM distinguishability between all combinations of ethnicities. Symbols depict specific ethnicities included in the ANOSIM tests, and boxes denote the R-value as a heatmap, where white indicates increasing and black indicates decreasing distinguishability relative to the R-value with all ethnicities. (B) Average ROC curves (for 10-fold cross-validation) and prediction performance metrics for one-versus-all RF classifiers for each ethnicity, using SMOTE (33) and down subsampling approaches for training.

Finally, to complement evaluation with ecological alpha and beta diversity we implement a random forest (RF) supervised learning algorithm to classify gut microbiota from genus level community profiles into their respective ethnicity. We build four one-versus-all binary classifiers to classify samples from each ethnicity compared to the rest, and use two different sampling approaches to train the models, Synthetic Minority Over-sampling Technique (SMOTE) (33) and down-sampling, for overcoming uneven representation of ethnicities in both the datasets (see Methods). Given that the area under the receiver operating characteristic (ROC) curve (or AUC) of a random guessing classifier is 0.5, the models classify each ethnicity fairly well (**Fig 3B**) with average AUCs across sampling techniques and datasets of 0.78 for Asian-Pacific Islanders, 0.76 for African Americans, 0.69 for Hispanics, and 0.70 for Caucasians. Ethnicity distinguishing RF taxa and out-of-bag error percentages appear in (**S2 Fig**).

### Recurrent taxon associations with ethnicity

Subtle to moderate ethnicity-associated differences in microbial communities may in part be driven by differential abundance of certain microbial taxa. 16.2% (130/802) of the AGP taxa and 20.6% (45/218) of HMP taxa across all classification levels (i.e. phylum to genus, **S5 Table**) significantly vary in abundance across ethnicities (Kruskal-Wallis, p_FDR_<0.05). Between datasets, 19.2% (25/130) of the AGP and 55.6% (25/45) of the HMP varying taxa replicate in the other dataset, representing a significantly greater degree of overlap than would be expected by chance (ethnic permutation analysis of overlap, p<0.001 each taxonomic level and all taxonomic levels combined). The highest replication of taxa varying by abundance occurs with 22.0% of families (9 significant in both datasets / 41 significantly varying families in either dataset), followed by genus with 13.4% (9 significant in both datasets / 67 significantly varying genera in either dataset).

Among 18 reproducible taxa, we categorize 12 as taxonomically distinct (**Fig 4**) and exclude 6 where nearly identical abundance profiles between family/genus taxonomy overlap. Comparing relative abundance differences between pairs of ethnicities for these 12 taxa in the AGP reveals 30 significant differences, of which 20 replicate in the HMP (p<0.05, Mann-Whitney-U). Intriguingly, all reproducible pairwise differences are a result of decreases in Asian-Pacific Islanders (**Fig 4**). We also test taxon abundance and presence/absence associations with ethnicity separately in the AGP using linear and logistic regression models respectively, and we repeat the analysis while incorporating categorical sex and continuous age and BMI as covariates (**S6 Table**). Clustering microbial families based on their abundance correlation reveals two co-occurrence clusters: (i) a distinct cluster of six Firmicutes and Tenericutes families in the HMP and (ii) an overlapping but more diverse cluster of 20 families in the AGP (**S3 Fig**). Nine of the 12 taxa found to recurrently vary in abundance across ethnicities are represented in these clusters (**Fig 4**), with four appearing in both clusters and the other five appearing either in or closely correlated with members of both clusters (**S3 Fig**). Furthermore, 90% (18/20) of families in the AGP cluster and 66% (4/6) of taxa in the HMP cluster significantly vary in abundance across ethnicities. We also found overlap for AGP and HMP datasets between taxa significantly varying in abundance across ethnicities (with FDR <0.05) and taxa in RF models with percentage importance greater than 50% for an ethnicity (**S2B Fig**). Taken together, these results establish general overlap of the most significant ethnicity-associated taxa between the these methods, reproducibility of microbial abundances that vary between ethnicities across datasets, and patterns of co-occurrence among these taxa which could suggest they are functionally linked.

**Fig 4.**
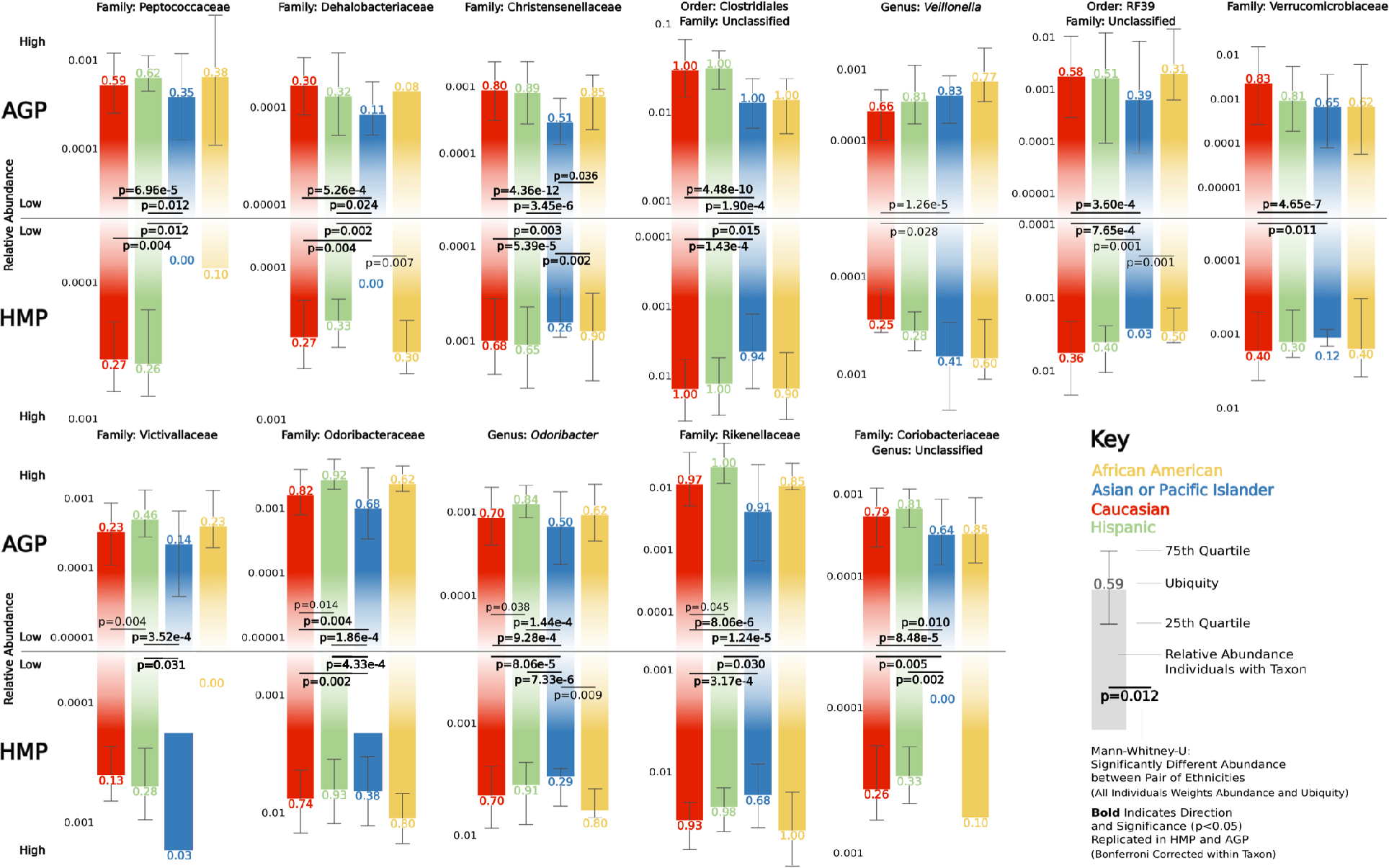
Ethnicity-associated taxa match between the HMP and AGP. Barplots depict the log10 transformed relative abundance for individuals possessing the respective taxon within each ethnicity, ubiquity appears above (AGP) or below (HMP) bars, and the 25^th^ and 75^th^ percentiles are shown with extending whiskers. Mann-Whitney-U tests evaluate differences in abundance and ubiquity for all individuals between pairs of ethnicities; for example, the direction of change in Victivallaceae is driven by ubiquity while abundance is higher for those possessing the taxon. Significance values are Bonferroni corrected for the six tests within each taxon and dataset, and bold p-values indicate that significance (p<0.05) and direction of change replicate in the AGP and HMP.

### Most heritable taxon of bacteria varies by ethnicity

Identified as the most heritable taxon in the human gut (34, 35), the family Christensenellaceae exhibits the second strongest significant difference in abundance across ethnicities in both AGP and HMP datasets (**S5 Table**, Family: AGP, Kruskal-Wallis, p_FDR_=1.55e-9; HMP, Kruskal-Wallis, p_FDR_=0.0019). Additionally, Christensenellaceae is variable by sex and BMI (AGP: Sex, Kruskal-Wallis, p_FDR_=1.22e-12; BMI, Kruskal-Wallis, p_FDR_=0.0020), and represents some of the strongest pairwise correlations with other taxa in both co-occurrence clusters (**S3 Fig**). There is at least an eight-fold and two-fold reduction in average Christensenellaceae abundance in Asian-Pacific Islanders relative to the other ethnicities in the AGP and HMP respectively (**S5 Table**), and significance of all pairwise comparisons in both datasets show reduced abundance in Asian-Pacific Islanders (**Fig 4**). Christensenellaceae also occur among the top 10 most influential taxa for distinguishing Asian-Pacific Islanders from other ethnicities using RF models for both AGP and HMP datasets (**S2A Fig**). Abundance in individuals possessing Christensenellaceae and presence/absence across all individuals significantly associate with ethnicity (**S6 Table**, Abundance, Linear Regression, p_Bonferroni_=0.006; Presence/Absence, Logistic Regression, p_Bonferroni_=8.802e-6), but there was only a slight correlation between the taxon’s relative abundance and BMI (**S4 Fig**). Confirming previous associations with lower BMI(36), we observe that AGP individuals with Christensenellaceae also have a lower BMI (Mean BMI, 23.7±4.3) than individuals without it (Mean BMI, 25.0±5.9; Mann-Whitney-U, p<0.001). This pattern is separately reflected in African Americans, Asian-Pacific Islanders, and Caucasians but not Hispanics (**Fig 5**), suggesting that each ethnicity may have different equilibria between the taxon’s abundance and body weight.

**Fig 5.**
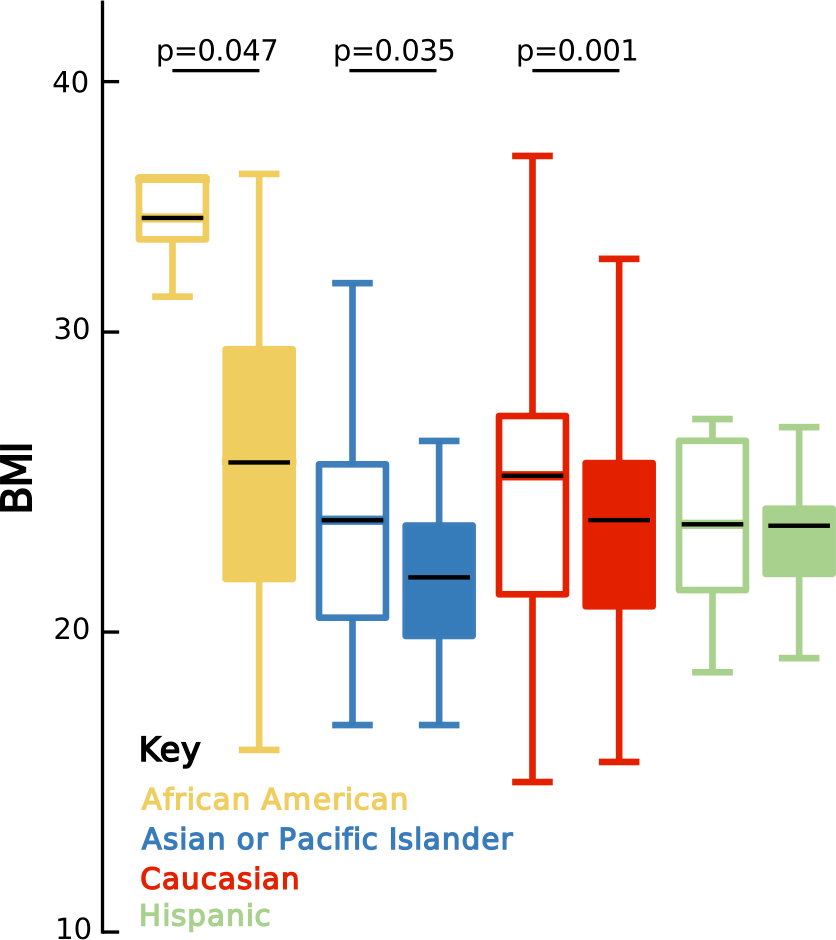
Christensenellaceae variably associate with BMI across ethnicities. Boxplots of BMI for individuals without (unfilled boxplots) and with (filled boxplots) Christensenellaceae. Significance was determined using one-tailed Mann-Whitney-U tests for lower continuous BMI values. Black lines indicate the mean relative abundance; the lower and upper end of each box represent the 25^th^ and 75^th^ quartiles respectively; and whiskers denote the 1.5 interquartile range.

### Genetic- and ethnicity-associated taxa overlap

Many factors associate with human ethnicity, including a small subset of population specific genetic variants (estimated ~0.5% genome wide) that vary by biogeographical ancestry (37, 38); self-declared ethnicity in the HMP is delineated by population genetic structure (20). Here we investigate whether ethnicity-associated taxa overlap with (i) taxa that have a significant population genetic heritability in humans (34, 35, 39, 40) and (ii) taxa linked with human genetic variants in two large Genome-Wide Association Studies (GWAS)-microbiota analyses (35, 40). All recurrent ethnicity-associated taxa except one were heritable in at least one study, with seven replicating in three or more studies (**Table 1**). Likewise, abundance differences in seven recurrent ethnicity-associated taxa demonstrate significant GWAS associations with at least one variant in the human genome. Therefore, we assess whether any genetic variants associated with differences in microbial abundance exhibit significant rates of differentiation (F_ST_) between 1,000 genomes superpopulations (38). Out of 49 variants associated with ethnically varying taxa, 21 have higher F_ST_ values between at least one pair of populations than that of 95% of other variants on the same chromosome and across the genome; the F_ST_ values of five variants associated with Clostridiaceae abundance rank above the top 99% (**S7 Table**). Since taxa that vary across ethnicities exhibit lower abundance in Asian-Pacific Islanders, it is notable that the F_ST_ values of 18 and 11 variant comparisons for East Asian and South Asian populations, respectively, are above that of the 95% rate of differentiation threshold from African, American, or European populations. Cautiously, the microbiota and 1,000 genomes datasets are not drawn from the same individuals, and disentangling the role of genetic from social and environmental factors will still require more controlled studies.

**Table 1.** Most recurrent ethnicity-associated taxa are previously reported heritable and genetically-associated taxa. The table shows population genetic heritability estimates and associated genetic variants for the 12 recurrent ethnically varying taxa. The minimum heritability cutoff was chosen as >0.1, and only exactly overlapping taxonomies were considered. Studies examined: ^A^UKTwins (2014, ‘A’ measure of additive heritability in ACE model) (34), ^B^Yatsunenko (2014, ‘A’ measure of additive heritability in ACE model) (34), ^C^UKTwins (2016, ‘A’ measure of additive heritability in ACE model) (35), ^D^Lim (2016, H2r measure of polygenic heritability in SOLAR (41)) (39), ^E^Turpin (2016, H2r measure of polygenic heritability in SOLAR (41)). *indicates excessive variants were excluded from table.

| Recurrent Ethnicity-Associated Taxa | Heritability | Genetic Associations |
| --- | --- | --- |
| Family: Peptococcaceae | 0.1213<sup>A</sup>, 0.2154<sup>C</sup>, 0.26<sup>E</sup> | rs143179968<sup>E</sup> |
| Family: Dehalobacteriaceae | 0.6878<sup>B</sup>, 0.3087<sup>C</sup> |  |
| Family: Christensenellaceae | 0.3819<sup>A</sup>, 0.6170<sup>B</sup>, 0.4230<sup>C</sup> |  |
| Order: Clostridiales, Family: Unclassified | 0.2914<sup>A</sup>, 0.4020<sup>B</sup>, 0.1330<sup>C</sup> | *40 Genetic Variants<sup>C</sup> |
| Genus: *Veillonella* | 0.1370<sup>A</sup>, 0.2168<sup>D</sup> | rs347941<sup>C</sup> |
| Order: RF39, Family: Unclassified | 0.2341<sup>A</sup>, 0.6618<sup>B</sup>, 0.3074<sup>C</sup> | rs4883972<sup>C</sup> |
| Family: Verrucomicrobiaceae | 0.1257<sup>A</sup>, 0.5973<sup>B</sup>, 0.1394<sup>C</sup> |  |
| Family: Victivallaceae |  |  |
| Family Odoribacteraceae | 0.1389<sup>A</sup>, 0.1917<sup>D</sup>, 0.34<sup>E</sup> | chr7:96414393<sup>E</sup>, rs115795847<sup>E</sup> |
| Genus: *Odoribacter* | 0.1916<sup>D</sup> |  |
| Family: Rikenellaceae | 0.1299<sup>D</sup>, 0.29<sup>E</sup> | rs17098734<sup>C</sup>, rs3909540<sup>C</sup>, rs147600757<sup>E</sup> |
| Family: Coriobacteraceae, Genus: Unclassified | 0.1364<sup>A</sup>, 0.2822<sup>B</sup>, 0.1609<sup>C</sup> | rs9357092<sup>E</sup> |

## Discussion

Many common diseases associate with microbiota composition and ethnicity, raising the central hypothesis that microbiota differences between ethnicities can occasionally serve as a mediator of health disparities. American’s self-declared ethnicity can capture socioeconomic, cultural, geographic, dietary and genetic diversity, and a similarly complex array of interindividual and environmental factors influence total microbiota composition. This complexity may result in challenges when attempting to recover consistent trends in total gut microbiota differences between ethnicities. The challenges in turn emphasize the importance of reproducibility, both through confirmation across analytical methods and replication across study populations (15–17, 20, 27, 42). In order to robustly substantiate the ethnicity-microbiota hypothesis, we evaluated recurrent associations between self-declared ethnicity and variation in both total gut microbiota and specific taxa in healthy individuals. Results provide hypotheses for examining specific members of the gut microbiota as mediators of health disparities.

Our findings from two American datasets demonstrate that: (i) ethnicity consistently captures gut microbiota with a slightly stronger effect size than other variables such as BMI, age, and sex, (ii) ethnicity is moderately predictable from total gut microbiota differences, and (iii) 12 taxa recurrently vary in abundance between the ethnicities, of which the majority have been previously shown to associate with human genetic variation. Whether shaped through socioeconomic, dietary, healthcare, genetic, or other ethnicity-related factors, reproducibly varying taxa represent sources for novel hypotheses addressing health disparities. For instance, the family Odoribacteraceae and genus *Odoribacter* are primary butyrate producers in the gut, and they have been negatively associated to severe forms of Crohn’s disease and Ulcerative Colitis in association with reduced butyrate metabolism (43–45). Asian-Pacific Islanders possess significantly less Odoribacteraceae and *Odoribacter* than Hispanics and Caucasians in both datasets, and severity of Ulcerative Colitis upon hospital admission has been shown to be significantly higher in Asian Americans (46). Considering broader physiological roles, several ethnicity-associated taxa are primary gut anaerobic fermenters and methanogens (47, 48), and associate with lower BMI and blood triglyceride levels (36, 49). Indeed, Christensenellaceae, Odoribacteraceae, *Odoribacter*, and the class Mollicutes containing RF39 negatively associate with metabolic syndrome and demonstrate significant population genetic heritability in twins (39). Implications for health outcomes warrant further investigation, but could be reflected by positive correlations of Odoribacteraceae, *Odoribacter*, Coriobacteriaceae, Christensenellaceae, and the dominant Verrucomicrobiaceae lineage *Akkermansia* with old age (50, 51). *Akkermansia* associations with health and ethnicity in western populations may reflect recently arising dietary and lifestyle effects on community composition, as this mucus consuming taxon is rarely observed in more traditional cultures globally (23). Moreover, these findings raise the importance of controlling for ethnicity in studies linking microbiota differences to disease because associations between specific microbes and a disease could be confounded by ethnicity of the study participants.

Based on correlations in individual taxon’s abundance, a similar pattern of co-occurrence previously identified as the ‘Christensenellaceae Consortium’ includes 11 of the 12 recurrent ethnically varying taxa (34), and members of this consortium associate with genetic variation in the human formate oxidation gene *ALDH1L1*, which is a genetic risk factor for stroke (35, 52, 53). Formate metabolism is a key step in the pathway reducing carbon dioxide to methane (54, 55), and increased methane associates with increased Rikenellaceae, Christensenellaceae, Odoribacteraceae and *Odoribacter* (56). Products of methanogenic fermentation pathways include short chain fatty acids such as butyrate, which through reduction of pro-inflammatory cytokines is linked to cancer cell apoptosis and reduced risk of colorectal cancer (57, 58). Asian Americans are the only ethnic group where cancer surpasses heart disease as the leading cause of death, and over 70% of Asian Americans were born overseas, which can affect assimilation into western lifestyles, leading to reduced access to healthcare and screening, and proper medical education (57, 59–61). Preliminary results from other groups suggest that the gut microbiome of Southeast Asian immigrants changes after migration to the United States (Dan Knights, personal communication). Indeed, as countries in Asia shift toward a more western lifestyle, the incidence of cancers, particularly gastrointestinal and colorectal cancers, are increasing rapidly, possibly indicating incompatibilities between traditionally harbored microbiota and western lifestyles (62–65). Asian Americans have higher rates of type 2 diabetes and pathogenic infections than Caucasians (66), and two metagenomic functions enriched in control versus type 2 diabetes cases appear to be largely conferred by cluster-associated butyrate-producing and motility-inducing Verrucomicrobiaceae and Clostridia taxa reduced in abundance among AGP and HMP Asian-Pacific Islanders (11). Both induction of cell motility and butyrate promotion of mucin integrity can protect against pathogenic colonization and associate with microbial community changes (11, 58, 67). Levels of cell motility and butyrate are key factors suspected to underlie a range of health disparities including inflammatory bowel disease, arthritis, and type 2 diabetes (11, 68–70). Patterns of ethnically varying taxa across ethnicities could result from many factors including varying diets, environmental exposures, sociocultural influences, human genetic variation and others. However, regardless of the mechanisms dictating assembly, these results suggest there is a reproducible, co-occurring group of taxa linked by similar metabolic processes known to promote homeostasis.

The utility of this work is establishing a framework for studying ethnicity-associated taxa and hypotheses of how changes in abundance or presence of these taxa may or may not shape health disparities, many of which also have genetic components. Differing in allele frequency across three population comparisons and associated with the abundance of Clostridiales, the genetic variant rs7587067 has a significantly higher frequency in African (Minor Allele Frequency (MAF)=0.802) versus East Asian (MAF=0.190, F_ST_=0.54, Chromosome=98.7%, Genome-Wide=98.9%), admixed American (MAF=0.278, F_ST_=0.44, Chromosome=99.0%, Genome-Wide=99.1%), and European populations (MAF=0.267, F_ST_=0.45, Chromosome=98.7.3%, Genome-Wide=98.7%). This intronic variant for the gene *HECW2* is a known eQTL (GTeX, eQTL Effect Size=−0.18, p=7.4e-5) (71, 72), and *HECW2* encodes a ubiquitin ligase linked to enteric gastrointestinal nervous system function through maintenance of endothelial lining of blood vessels (73, 74). Knockout of *HECW2* in mice reduced enteric neuron networks and gut motility, and patients with Hirschsprung’s disease have diminished localization of *HECW2* to regions affected by loss of neurons and colon blockage when compared to other regions of their own colon and healthy individuals (75). Hirschsprung’s disease presenting as full colon blockage is rare and has not undergone targeted examination as a health disparity, however a possible hypothesis is that lower penetrance of the disease in individuals with the risk allele at rs7587067 could lead to subtler effects on gut motility resulting in Clostridiales abundance differences.

Despite the intrigue of connecting the human genome, microbiota and disease phenotypes, evaluating such hypotheses will require more holistic approaches including incorporating metagenomics and metabolomics to identify whether enzymes or metabolic functions reproducibly vary across ethnicities, as well as direct functional studies in model systems to understand if correlation is truly driven by causation. Further limitations should also be considered, including recruitment biases for the AGP versus HMP, variation in sample processing and OTU clustering, and uneven sampling which could only be addressed with down sampling of over-represented ethnicities. Still, despite these confounders care was taken to demonstrate the reproducibility of results across statistical methods, ecological metrics, rarefaction depths, and study populations. Summarily, this work suggests that abundance differences of specific taxa, rather than whole communities, may represent the most reliable ethnic signatures in the gut microbiota. A reproducible co-occurring subset of these taxa link to a variety of overlapping metabolic processes and health disparities, and contain the most reproducibly heritable taxon, Christensenellaceae. Moreover, a majority of the microbial taxa associated with ethnicity are also heritable and genetically-associated taxa, suggesting there is a possible connection between ethnicity and genetic patterns of biogeographical ancestry that may play a role in shaping these taxa. Our results emphasize the importance of sampling ethnically diverse populations of healthy individuals in order to discover and replicate ethnicity signatures in the human gut microbiota, and they highlight a need to account for ethnic variation as a potential confounding factor in studies linking microbiota differences to disease. Further reinforcement of these results may lead to generalizations about microbiota assembly and even consideration of specific taxa as potential mediators or treatments of health disparities.

## Materials and Methods

### Data Acquisition

AGP data was obtained from the project FTP repository located at *ftp://ftp.microbio.me/AmericanGut/*. AGP data generation and processing prior to analysis can be found at: *https://github.com/biocore/American-Gut/tree/master/ipynb/primary-processing*. All analyses utilized the rounds-1-25 dataset which was released on March 4, 2016. Throughout all analyses, QIIME v1.9.0 was used in an Anaconda environment [*https://continuum.io*] for all script calls, custom scripts and notebooks were run in the QIIME 2 Anaconda environment with python version 3.5.2, and plots were post-processed using Inkscape [*https://inkscape.org/en/*] (76). Ethnicity used in this study was self-declared by AGP study participants as one of four groups: African American, Asian or Pacific Islander (Asian-Pacific Islander), Caucasian, or Hispanic. Sex was self-declared as either male, female, or other. Age was self-declared as a continuous integer of years old, and age categories defined by the AGP by decade (i.e. 20’s, 30’s…) were used in this study. BMI was self-declared as an integer, and BMI categories defined by AGP of underweight, healthy, overweight, and obese were utilized. A total of 31 categorical metadata factors were assessed for structuring across ethnicities with a two proportion Z test between pairs of ethnicities using a custom python script (**S1 Table** additional sheets). The p-values were Bonferroni corrected within each metadata factor for the number of pairwise ethnic comparisons. 97% Operational Taxonomic Units (OTUs) generated for each dataset are utilized throughout to maintain consistency with other published literature, however microbial taxonomy of the HMP is reassigned using the Greengenes reference database (77). Communities characterized with 16S rDNA sequencing of variable region four followed an identical processing pipeline for all samples, which was developed and optimized for the Earth Microbiome Project (78). HMP 16S rDNA data processed using QIIME for variable regions 3-5 was obtained from *http://hmpdacc.org/HMQCP/*. Demographic info for individual HMP participants was obtained through dbGaP restricted access to study phs000228.v2.p1, with dbGaP approval granted to SRB and non-human subjects determination IRB161231 granted by Vanderbilt University. Ethnicity and sex were assigned to subjects based on self-declared values, with individuals selecting multiple ethnicities being removed unless they primarily responded as Hispanic, while categorical age and BMI were established from continuous values using the same criteria for assignment as in AGP. The HMP Amerindian population was removed due to severe under-representation. This filtered HMP table was used for community level analyses (ANOSIM, Alpha Diversity, beta intra-inter), however to allow comparison with the AGP dataset, community subset analyses (co-occurrence, abundance correlation, etc…) were performed with taxonomic assignments in QIIME using the UCLUST method with the GreenGenes_13_5 reference.

### Quality Control

AGP quality control was performed in Stata v12 (StataCorp, 2011) using available metadata to remove samples (Raw N=9,475): with BMI more than 60 (−988 [8,487]) or less than 10 (−68 [8,419]), missing age (−661 [7,758]), with age greater than 55 years old (−2,777 [4,981]) or less than 18 years old (−582 [4,399]), and blank samples or those not appearing in the mapping file (−482 [3,917]), with unknown ethnicity or declared as other (−131 [3786]), not declared as a fecal origin (−2,002 [1784]), with unknown sex or declared as other (−98 [1686]), or located outside of the United States (−209 [1477]). No HMP individuals were missing key metadata or had other reasons for exclusion (−0[298]). Final community quality control for both AGP and HMP was performed by filtering OTUs with less than 10 sequences and removing samples with less than 1,000 sequences (AGP, −102 [1375]; HMP, −0 [298]). All analyses used 97% OTUs generated by the AGP or HMP, and unless otherwise noted, results represent Bray-Curtis beta diversity and Shannon alpha diversity at a rarefaction depth of 1,000 counts per sample.

### ANOSIM, PERMANOVA, and BioEnv Distinguishability

The ANOSIM test was performed with 9,999 repetitions on each rarefied table within a respective rarefaction depth and beta diversity metric (**Fig 1** & **S2A-B Table**), with R-values and p-values averaged across the rarefactions. Consensus beta diversity matrices were calculated as the average distances across the 100 rarefied matrices for each beta diversity metric and depth. Consensus distance matrices were randomly subsampled ten times for subset number of individuals from each ethnic group with more than that subset number prior to ANOSIM analysis with 9,999 repetitions, and the results were averaged evaluating the effects of more even representations for each ethnicity (**S2C Table**). Consensus distance matrices had each ethnicity and pair of ethnicities removed prior to ANOSIM analysis with 9,999 repetitions, evaluating the distinguishability conferred by inclusion of each ethnicity (**Fig 3A**, **S2F Table**). Significance was not corrected for the number of tests to allow comparisons between results of different analyses, metrics, and depths. PERMANOVA analyses were run using the R language implementation in the Vegan package (79), with data handled in a custom R script using the Phyloseq package (80). Categorical variables were used to evaluate the PERMANOVA equation (Beta-Diversity Distance Matrix ~ Ethnicity + Age + Sex + BMI) using 999 permutations to evaluate significance, and the R and p values were averaged across 10 rarefactions (**S2D Table**). The BioEnv test, or BEST test, was adapted to allow evaluation of the correlation and significance between beta diversity distance matrices and age, sex, BMI, and ethnicity simultaneously (**S2E Table**) (29). At each rarefaction depth and beta diversity metric the consensus distance matrix was evaluated for its correlation with the centered and scaled Euclidian distance matrix of individuals continuous age and BMI, and categorical ethnicity and sex encoded using patsy (same methodology as original test)[*https://patsy.readthedocs.io/en/latest/#*]. The test was adapted to calculate significance for a variable of interest by comparing how often the degree of correlation with all metadata variables (age, sex, BMI, ethnicity) was higher than the correlation when the variable of interest was randomly shuffled between samples 1,000 times.

### Alpha Diversity

Alpha diversity metrics (Shannon, Simpson, Equitability, Chao1, Observed OTUs) were computed for each rarefied table (QIIME: alpha_diversity.py), and results were collated and averaged for each sample across the tables (QIIME: collate_alpha.py). Pairwise nonparametric t-tests using Monte Carlo permutations evaluated alpha diversity differences between the ethnicities with Bonferroni correction for the number of comparisons (**Fig 2A**, **S3 Table**, QIIME: compare_alpha_diversity.py). A Kruskal-Wallis test implemented in python was used to detect significant differences across all ethnicities.

### Beta Diversity

Each consensus beta diversity distance matrix had distances organized based on whether they represented individuals of the same ethnic group, or were between individuals of different ethnic groups. All values indicate that all pairwise distances between all individuals were used (**Fig 2B**, **S4A-B Table**), mean values indicate that for each individual their average distance to all individuals in the comparison group was used as a single point to assess pseudo-inflation (**S4C-D Table**). A Kruskal-Wallis test was used to calculate significant differences in intra-ethnic distances across all ethnicities. Pairwise Mann-Whitney-U tests were calculated between each pair of intra-ethnic distance comparisons, along with intra-versus-inter ethnic distance comparisons. Significance was Bonferroni corrected within the number of intra-intra-ethnic and intra-inter-ethnic distance groups compared, with violin plots of intra- and inter-ethnic beta diversity distances generated for each comparison.

### Random Forest

RF models were implemented using taxa summarized at genus level, which performed better compared to RF models using OTUs as features, both in terms of classification accuracy and computational time. We first rarefied OTU tables at sequence depth of 10,000 (using R v3.3.3 package *vegan’s* rrarefy() function) and then summarized rarefied OTUs at genus-level (or lower characterized level if genus was uncharacterized for an OTU). We filtered for rare taxa by removing taxa present in fewer than half of the number of samples in rarest ethnicity (i.e. fewer than 10/2 = 5 samples in HMP and 13/2 = 6 (rounded down) in AGP), retaining 85 distinct taxa in HMP dataset and 322 distinct taxa in AGP dataset at genus level. The resulting taxa were normalized to relative abundance and arcsin-sqrt transformed before being used as features for the RF models. We initially built multi-class RF model, but since the RF model is highly sensitive to the uneven representation of classes, all samples were identified as the majority class, i.e. Caucasian. In order to even out the class imbalance, we considered some sampling approaches, but most existing techniques for improving classification performance on imbalanced datasets are designed for binary class imbalanced datasets, and are not effective on datasets with multiple underrepresented classes. Hence, we adopted the binary classification approach and built four one-versus-all binary RF classifiers to classify samples from each ethnicity compared to the rest. 10-fold cross-validation (using R package *caret* (81)) was performed using ROC as the metric for selecting optimal model. The performance metrics and ROC curves were averaged across the 10 folds (**Fig 3B**). Without any sampling during training the classifiers, most samples were identified as the majority class, i.e. the Caucasian, by all four one-versus-all RF classifiers. In order to overcome this imbalance in class representation, we applied two sampling techniques inside cross-validation: i) down-sampling, and ii) Synthetic Minority Over-sampling Technique (or SMOTE) (33). In the down-sampling approach, the majority class is down-sampled by random removal of instances from the majority class. In the SMOTE approach, the majority class is down-sampled and synthetic samples from the minority class are generated based on k-nearest neighbors technique (33). Note, the sampling was performed inside cross-validation on training set, while the test was performed on unbalanced held-out test set in each fold. In comparison to a no-sampling approach, which classified most samples as the majority class, i.e., Caucasians, our sampling-based approach leads to improved sensitivity for classification of minority classes on unbalanced test sets. Nevertheless, the most accurate prediction remains for the inclusion in the majority class.

The ROC curves and performance metrics table in **Fig 3B** show the sensitivity-specificity tradeoff and classification performance for one-versus-all classifier for each ethnicity for both the sampling techniques applied on both the datasets. For both the datasets, down-sampling shows higher sensitivity and lower specificity and precision for minority classes (i.e. African Americans, Asian-Pacific Islanders and Hispanics) compared to SMOTE. However, for the majority class (i.e. Caucasian), down-sampling lowers the sensitivity and increases the specificity and precision compared to SMOTE. The sensitivity-specificity tradeoff, denoted by the area under the ROC curve (or AUC) is reduced for Hispanics in both the datasets. The most important taxa with >50% importance for predicting an ethnicity using RF model with SMOTE sampling approach are shown in **S2A Fig**. Among the 10 most important taxa for each ethnicity, there are 9 taxa which overlap between the AGP and HMP datasets (highlighted by the blue rectangular box); however, which ethnicity they best distinguish varies between the two datasets. Within each dataset we highlighted taxa which are distinguishing in RF models and have distinguishing differential abundance in **S2B Fig**, reporting both the FDR corrected significance for Kruskal-Wallis tests of differential abundance, and the percent importance for the most distinguished ethnicity of each in RF models. We also report out-of-bag errors for the final RF classifier that was built using the optimal model parameters obtained from cross-validation approach corresponding to each ethnicity and sampling procedure for both AGP and HMP datasets in **S2C Fig**.

### Taxon Associations

Taxon differential abundance across categorical metadata groups was performed in QIIME (QIIME: group_significance.py, **S5 Table**) to examine whether observation counts (i.e. OTUs and microbial taxon) are significantly different between groups within a metadata category (i.e. ethnicity, sex, BMI, age). The OTU table prior to final community quality control was collapsed at each taxonomic level (i.e. Phylum – Genus; QIIME: collapse_taxonomy.py), with counts representing the relative abundance of each microbial taxon. Differences in the mean abundance of taxa between ethnicities were calculated using Kruskal-Wallis nonparametric statistical tests. P-values are provided alongside false discovery rate and Bonferroni corrected P-values, and taxon were ranked from most to least significant. Results were collated into excel tables by taxonomic level and metadata category being examined, with significant (false discovery rate and Bonferroni P-value < 0.05) highlighted in orange, and taxa that were false discovery rate significant in both datasets were colored red. The Fisher’s exact test for the overlap of number of significant taxa between datasets was run at the online portal (http://vassarstats.net/tab2×2.html), with the expected overlap calculated as 5% of the number of significant taxa at all levels within the respective dataset, and the observed 25 taxa that overlapped in our analysis. The permutation analysis was performed by comparing the number of significant taxa (**S5 Table**, p_FDR_<0.05) overlapping between the AGP and HMP to the number overlapping when the Kruskal-Wallis test was performed 1,000 times with ethnicity randomly permuted. In 1/1000 runs there was one significant taxon overlapping at the family level, and one in 3/1000 permutations at the genus level, with no significant taxa overlapping in any repetitions at higher taxonomic levels. The 12 families and genera that were significantly different were evaluated to not be taxonomically distinct if their abundances across ethnicities at each level represented at least 82-100% (nearly all >95%) of the overlapping taxonomic level, and the genera was used if classified, and family level used if genera was unclassified (g_). Average relative abundances on a log10 scale among individuals possessing the taxon were extracted for each taxon within each ethnicity, and the abundance for 12 families and genera were made into barchart figures (**Fig 4**). The external whisker (AGP above, HMP below) depict the 75^th^ quartile of abundance, and the internal whisker depicts the 25^th^ quartile. Pairwise Mann-Whitney-U tests were performed between each pair of ethnicities using microbial abundances among all individuals, and were Bonferroni corrected for the six comparisons within each taxon and dataset. Bonferroni significant P-values are shown in the figure, and shown in bold if significance and direction of change replicate in both datasets. Ubiquity shown above or below each bar was calculated as the number of individuals in which that taxon was detected within the respective ethnicity. Additional confirmation of ethnically varying abundance was also performed at each taxonomic level (**S6 Table**), where the correlation of continuous age and BMI along with categorically coded sex and ethnicity were simultaneously measured against the log 10 transformed relative abundance of each taxon among individuals possessing it using linear regression (**S6 Table** - Abundance), and against the presence or absence of the taxon in all individuals with logistic regression (**S6 Table** - Presence Absence). Significance is presented for the models each with ethnicity alone, and with all metadata factors included (age, sex, BMI), alongside Bonferroni corrected p-values, and individual effects of each metadata factor.

### Co-Occurrence Analysis

Bacterial taxonomy was collapsed at the family level, Spearman correlation was calculated between each pair of families using SciPy (82), and clustermaps were generated using seaborn (**S3 Fig**), and ethnic associations were drawn from **S5 Table**. Correlations were masked where Bonferroni corrected Spearman p-values were >0.05, and clusters were identified as the most prominent (strongest correlations) and abundance enriched. Enrichment of ethnic association was evaluated by measuring the Mann-Whitney-U of cluster families ethnic associations (p-values, **S5 Table**) compared to the ethnic associations of non-cluster taxa. Cluster associated families were identified as having at least three significant correlations with families within the cluster.

### Christensenellaceae Analysis

The abundance of the family Christensenellaceae was input as relative abundance across all individuals from the family level taxonomic table. Individuals were subset based on the presence/absence of Christensenellaceae and BMIs were compared using a one tailed Mann-Whitney-U test, then each was further subset by ethnicity and BMI compared using one tailed Mann-Whitney-U tests and boxplots within each ethnicity (**Fig 5**).

### Genetically Associated, Heritable, and Correlated Taxa Analysis

Genetically associated taxa from population heritability studies (34, 35, 39, 40) with a minimum heritability (A in ACE models or H2r) >0.1, and from GWAS studies (35, 40) were examined for exact taxonomic overlap with our 12 ethnically-associated taxa. The 42 genetic variants associated with Unclassified Clostridiales are: rs16845116, rs586749, rs7527642, rs10221827, rs5754822, rs4968435, rs17170765, rs1760889, rs6933411, rs2830259, rs7318523, rs17763551, rs2248020, rs1278911, rs185902, rs2505338, rs6999713, rs5997791, rs7236263, rs10484857, rs9938742, rs1125819, rs4699323, rs641527, rs7302174, rs2007084, rs2293702, rs9350764, rs2170226, rs2273623, rs9321334, rs6542797, rs9397927, rs2269706, rs4717021, rs7499858, rs10148020, rs7524581, rs11733214, rs7587067 from (35). These 40 variants along with variants in **Table 1** except for chr7:96414393 (total=49) were then assessed in 1,000 Genomes individuals for significant differentiation across superpopulations (38). The 1,000 Genomes VCF files were downloaded (ftp://ftp.1000genomes.ebi.ac.uk/vol1/ftp/release/20130502/), and variants with a minor allele frequency less than 0.01 were removed with F_ST_ calculated between each pair of superpopulations using vcftools (83). The East Asian versus South Asian F_ST_ rates were not used in the analysis. A custom script was used to examine the F_ST_ for each of the 49 variants and compare to the F_ST_ of all variants on the same chromosome and all variants genome-wide for that pair of populations, with percentile calculated and the number of variants with a higher F_ST_ divided by the total number of variants. The eQTL value and significance for rs7587067 were drawn from the GTEx database (72).

## Data and Code Availability

Code, scripts, and data underlying figures are publicly available from the GitHub repository [https://github.com/awbrooks19/microbiota_and_ethnicity]. Individual metadata (age, sex, ethnicity…) for the Human Microbiome Project are held under restricted access available through dbGaP application [NCBI - dbGaP, Human Microbiome Project, https://www.ncbi.nlm.nih.gov/projects/gap/cgi-bin/study.cgi?study_id=phs000228.v3.p1].

## Acknowledgements

This work was supported by National Institutes of Health training grants 4T32GM08017810, 5T32GM08017809, and 5T32GM0817808 to AWB, the Vanderbilt Office of Equity, Diversity and Inclusion to A.W.B. and S.R.B., the Vanderbilt Microbiome Initiative to S.R.B., and the Alfred P. Sloan Foundation Fellowship to R.B‥ The content is solely the responsibility of the authors and does not necessarily represent the official views of the National Institutes of Health. We thank the American Society of Microbiology for supporting travel to present this work. We would also like to thank Tony Capra, David Samuels, Patrick Abbot, Antonis Rokas, and other members of the Vanderbilt Genetics Institute and Bordenstein Lab for input. The authors acknowledge the Minnesota Supercomputing Institute (MSI) at the University of Minnesota and the Advanced Computing Center for Research and Education (ACCRE) at Vanderbilt University for providing resources that contributed to the research results reported within this paper.

## Contributions

A.W.B., S.P., R.B., and S.R.B. conceived and designed the research. A.W.B. performed, analyzed, and interpreted all experiments with the exception of the RF analysis planned and performed by S.P. and R.B. S.R.B. supervised all experimental designs, data analysis, and data interpretation. All authors participated in manuscript preparation, editing, and final approval.

## Supplementary Table/Figure Legends

**S1 Fig.**
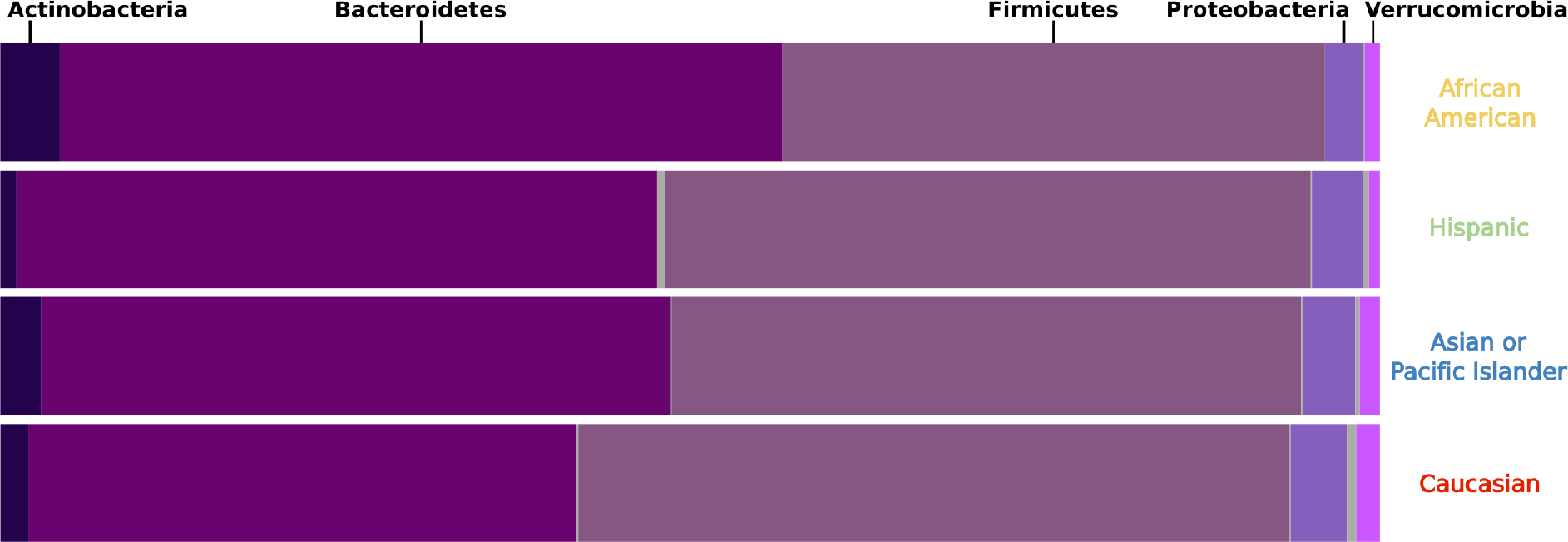
The average relative abundance of dominant microbial phyla for each ethnicity.

**S2 Fig.**
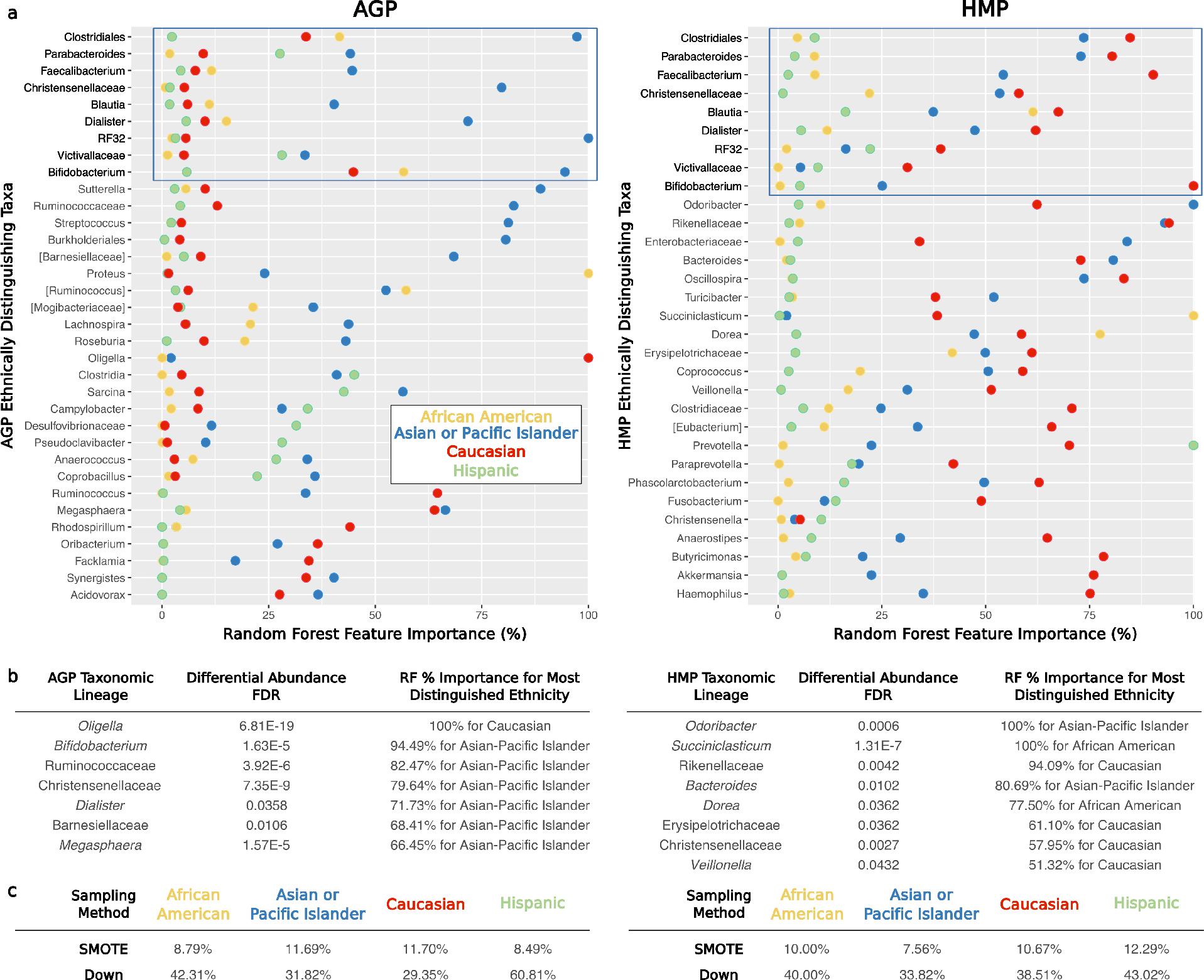
Summary of RF distinguishing taxa and out-of-bag error for each ethnicity. (A) Importance of taxa for predicting each ethnicity using RF models with SMOTE sampling approach are shown as percentage contributions, highlighted by color for each ethnicity. Among the 10 most important taxa for each ethnicity, 9 overlap between the AGP and HMP datasets (highlighted by the blue rectangular box), however which ethnicity they best distinguish varies between the two datasets. (B) Taxa which are distinguishing in RF models and have distinguishing differential abundance in **S5 Table**. The FDR corrected significance for Kruskal-Wallis tests of differential abundance and the percent importance for the most distinguished ethnicity of each in RF models are shown. (C) Out-of-bag error percentages for the final RF classifier that was built using the optimal model parameters obtained from cross-validation approach corresponding to each ethnicity and sampling procedure for both AGP and HMP datasets.

**S3 Fig.**
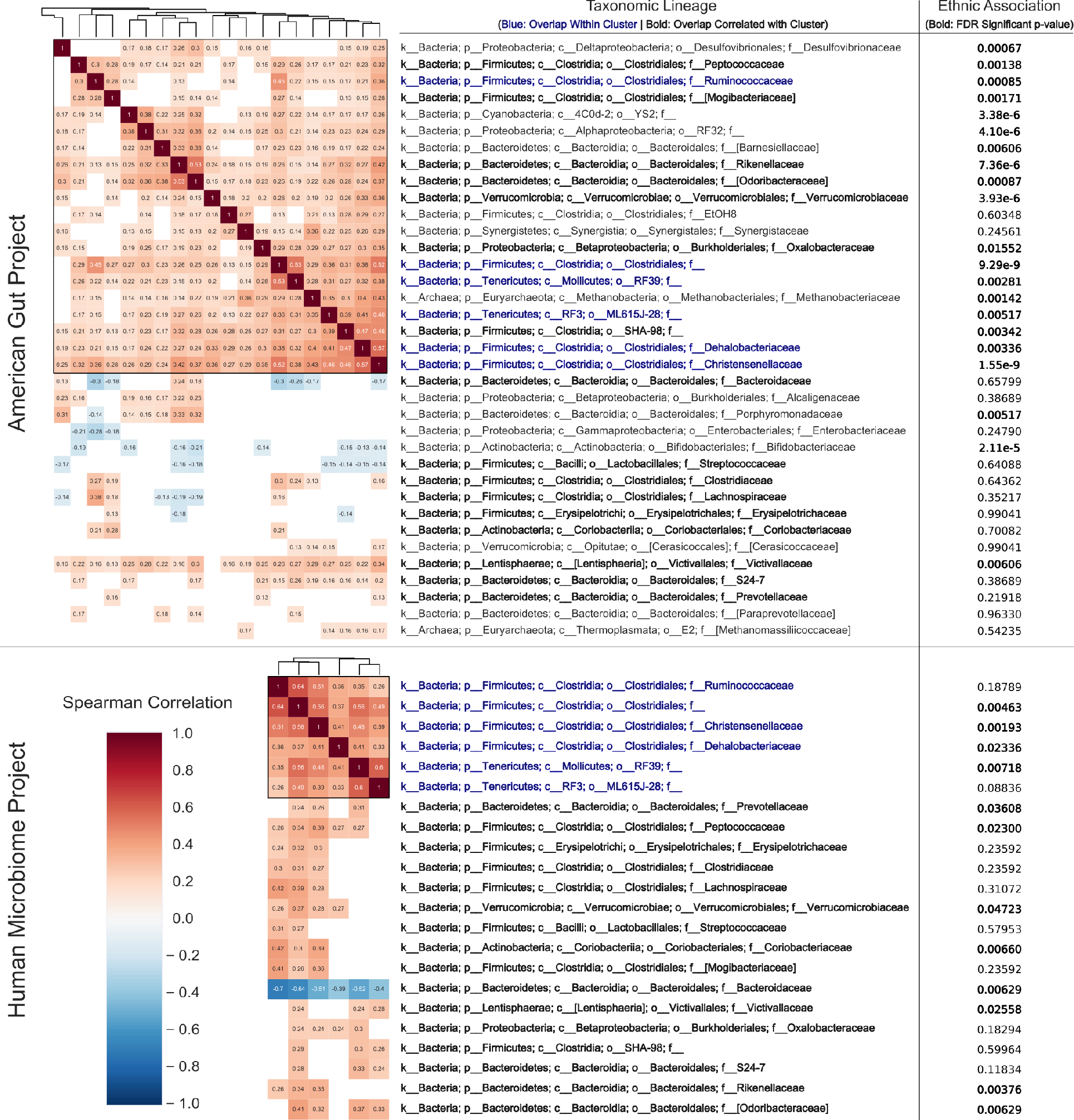
Abundance correlation of microbial families. Spearman correlation clustermaps of bacterial abundance for families in the AGP and HMP. Numbers within boxes depict the spearman correlation value with heatmap coloration from blue negative correlation (−1), white no correlation (0), to red positive correlation (1). Positions have been masked based on Bonferroni significance <0.05 for the total clustermap of all microbial families. Taxa within boxes were identified as a highly correlated cluster, and taxa outside the boxes share multiple correlations with those within the cluster. Blue taxonomic names indicate overlap of taxa within boxes of both the AGP and HMP, while black indicate multiple correlations with the clusters in both datasets. The ethnic association column depicts FDR corrected p-values from Kruskal-Wallis tests in **S5 Table**, which are bolded if <0.05.

**S4 Fig.**
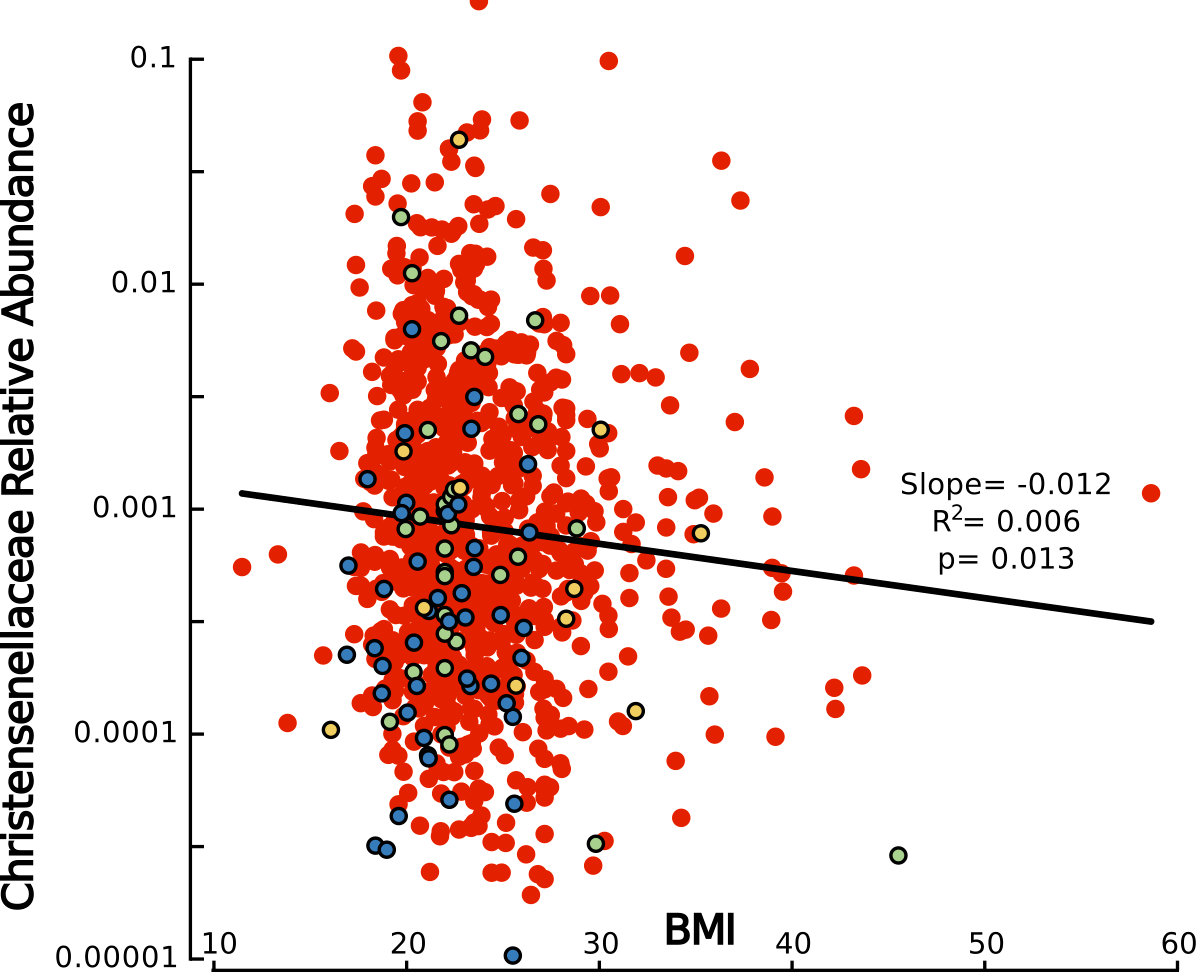
Correlation of BMI with Christensenellaceae abundance. The relationship for each individual between log10 transformed Christensenellaceae abundance on the y axis and BMI on the x axis, with statistics slope, R^2^, and p fit with a linear regression. Coloration of each point indicates ethnicity: Yellow – African American; Blue – Asian-Pacific Islander; Green – Hispanic; Red – Caucasian.

**S1 Table. Demographic information for the AGP.** Breakdown of age and BMI by sex and ethnicity. Heatmaps were constructed within each statistic and category (bounded by black box). The means for all sex and ethnic groups were used as the center (white), with higher values indicated in red and lower in blue. HMP data is not shown because of data access restrictions on participant metadata, available through dbGaP application. Additional sheets depict proportions tests of ethnic structuring for 31 metadata factors, each on their own sheet.

**S2 Table. Microbiota distinguishability by ethnicity, age, sex and BMI.** (A) AGP and HMP ANOSIM distinguishability by ethnicity, age, sex, and BMI at a rarefaction depth of 1,000 and across four ecological metrics (more details in table). (B) AGP ANOSIM distinguishability by ethnicity, age, sex, and BMI at rarefaction depths of 1,000 and 10,000. (C) ANOSIM results for consensus distance matrix while subsampling the maximum number of individuals from each ethnic group. (D) BioEnv results of correlation between ethnicity, age, sex, and BMI together with outcome as multivariate beta diversity distance matrices [Distance Matrix = Ethnicity*x1 + Categorical Age*x2 + Categorical BMI*x3 + Sex*x4 + B]. (E) ANOSIM results for consensus distance matrix when each ethnicity and group of ethnicities are sequentially removed from the analysis.

**S3 Table. Alpha diversity by ethnicity, age, sex and BMI.** Alpha Diversity for Ethnicity, Age, Sex, and BMI across varying rarefaction depths and beta diversity metrics in AG (4A, 4C-E), and for ethnicity in the HMP (4B). Results are based on non-parametric permutation based t-tests, and p-values are Bonferroni corrected within each factor of interest, depth, and metric.

**S4 Table. Comparison of beta diversity distances for within and between ethnicities.** All values depicted are Mann-Whitney-U p-values. (A) All distances between pairs of individuals within each ethnicity were compared between ethnicities across rarefaction depths 1,000 and 10,000, four beta diversity metrics, and with while subsampling over-represented ethnicities. (B) All distances between pairs of individuals within and between each ethnicity were compared between ethnicities. (C) Mean distances between pairs of individuals within each ethnicity were compared between ethnicities. (D) Mean distances between pairs of individuals within and between each ethnicity were compared between ethnicities.

**S5 Table. Taxa which are differentially abundant by ethnicity, sex, BMI, and age in the AGP and HMP.** Kruskal-Wallis results for differential taxa abundance across metadata groupings, including FDR and Bonferroni corrected p-values, and taxa abundance averages within each group. Metadata factors and taxonomic levels are separated by excel tabs.

**S6 Table. Taxa which are correlated with ethnicity, sex, BMI, and age in the AGP.** Results of linear (Abundance) and logistic (Presence Absence) regression results for differential taxa abundance across metadata factors separated by taxonomic level. Columns in order indicate the taxon name, the number of individuals with non-zero abundance; then the p-value for ethnicity alone, the p-value Bonferroni corrected, the f-test statistic, and R^2^; then the same values for the regression with ethnicity, age, sex, and BMI together; then the abundances in each ethnic group, and finally the p-values for each factor broken down.

**S7 Table. Genetic variants with taxa associations and detailed 1,000 Genomes population differentiation rates (F_ST_).** Variants in red indicate the variant has at least one F_ST_ above the 95^th^ percentile for high differentiation between at least one pair of populations. Columns I-BU represent the values for calculating variant F_ST_ and percentiles. The first two spaces indicate the two superpopulations being compared. F_ST_ indicates the rate of differentiation for that variant between that pair of populations. Higher indicates the number of variants genome-wide with a higher F_ST_, and total indicates the total genome-wide variants examined. The columns with chromosome indicate the number of variants with higher F_ST_ and total variants on the same chromosome as the variant of interest. Percent indicates the number of variants with a higher F_ST_ divided by the total number of variants.

## Abbreviations

AGP: American Gut Project
ANOSIM: Analysis of Similarity
AUC: Area Under the Curve
BMI: Body Mass Index
F_ST_: Fixation Index
GWAS: Genome-Wide Association Studies
HMP: Human Microbiome Project
MAF: Minor Allele Frequency
OTU: Operational Taxonomic Unit
PERMANOVA: Permutational Multivariate Analysis of Variance
RF: Random Forest
ROC: Receiver Operating Characteristic
SMOTE: Synthetic Minority Over-sampling Technique

